# Biological changes, political ideology, and scientific communication shape human perceptions of pollen seasons

**DOI:** 10.1101/2024.07.30.605525

**Authors:** Yiluan Song, Adam Millard-Ball, Nathan Fox, Derek Van Berkel, Arun Agrawal, Kai Zhu

## Abstract

Climate change is changing the timing and intensity of pollen seasons, thereby increasing human exposure to allergenic pollen. Climate-driven changes in pollen seasons present a unique opportunity to craft messaging aimed at communicating climate change impacts on public health through changes in the biological systems. However, it is unclear how pollen seasons are experienced and understood by the public, including how well we detect pollen seasons or what factors we view as responsible for changes in pollen seasons. Here, we use social media data in the United States from 2012 to 2022 to assess public perceptions of pollen seasons at large spatiotemporal scales. We show that pollen seasons detected by social media (Twitter) users align well with natural pollen seasons. Attribution of changing pollen seasons, however, varies based on political ideology: liberal users, compared to conservative users, are more likely to attribute changing pollen seasons to climate change. Mass media and scientific experts play a key role in communicating how climate change drives changing pollen seasons. Our findings reveal how changes in biological systems, political ideology, and scientific communications collectively shape public perceptions of pollen seasons under climate change. Our findings constitute a step towards more effective climate change impact communication and public health intervention design.

## 1. Introduction

Changes in phenology, the timing of recurring biological events, are among the most sensitive biological responses to climate change (Parmesan & Yohe, 2003). In particular, climate change has shifted pollen phenology, the timing and intensity of plant pollen production, release, and dispersal, leading to generally earlier starts (at a rate of up to 7 days decade^-1^), longer durations (up to 11 days decade^-1^) (Anderegg et al., 2021; van Vliet et al., 2002; Y. Zhang et al., 2015; Y. Zhang & Steiner, 2022; Ziska et al., 2011), and greater intensities (Anderegg et al., 2021; Ziska et al., 2019) of pollen seasons. Climate change is therefore anticipated to increase human exposure to pollen, elevating the risks of allergic asthma and allergic rhinitis (hay fever) (D’Amato et al., 2014; Lake et al., 2017; Sapkota et al., 2020; Traidl-Hoffmann et al., 2003). In addition to allergies, pollen shapes diverse human experiences: observations of pollen can mark the coming of pleasant spring weather, while pollen cover on cars is often seen as a nuisance. Because of the potentially profound impacts on the public, the changing pollen seasons may serve to frame effective communication of climate change impacts for climate change mitigation and adaptation.

Climate-driven changes in pollen seasons provide a promising yet understudied narrative (a “storyline”) in the framing of climate change. A frame around changes in the biological system has generally been effective in improving climate change engagement, with a risk of distancing people from the impacts (Li & Su, 2018; Maibach et al., 2010). An emerging frame around public health has the potential to make the problem of climate change understandable and personally meaningful, but it has not always been effective in practice (Li & Su, 2018; Maibach et al., 2010). At the intersection of the biological frame and the public health frame, a narrative of climate-driven changes in pollen seasons may have a unique advantage, given how personally and broadly it is experienced . Nevertheless, this narrative shares two potential bottlenecks with some other narratives (Moser & Ekstrom, 2010). The first bottleneck lies in the detection of changes in the environment. The public relies heavily on heuristics when noticing, reporting, and recalling physical changes in the environment (Howe & Leiserowitz, 2013; Kirilenko et al., 2015; Moore et al., 2019), such that individuals’ perceptions of pollen phenology may not match natural processes. Previous studies have suggested alignment between airborne pollen concentration and the count of related social media posts (Gesualdo et al., 2015; K. Lee et al., 2015; Cowie et al., 2018; Wakamiya et al., 2019) at small spatiotemporal scales (local or intra-annual). In cases where people do accurately detect such environmental change, this may not be attributed to climate change, leading to an additional bottleneck. Such attribution processes are influenced by psychological, demographic, cultural, and political factors (Akerlof et al., 2013; Hulme, 2014; Lahsen & Ribot, 2022; Myers et al., 2013; Reser & Bradley, 2020; van der Linden, 2015). In particular, ideological orientations affect climate change attribution (Dunlap et al., 2016; Farrell, 2016; Guilbeault et al., 2018). For example, those with liberal values are more likely to associate the occurrence of extreme weather events with climate change (Cutler, 2015; Howe & Leiserowitz, 2013). More than an individual-level process, climate change attribution might be influenced by scientific communication. In this study, scientific communication specifically refers to the dissemination of scientific knowledge by mass media (Carmichael & Brulle, 2017; Egan & Mullin, 2017) and scientific experts (Anderegg et al., 2010).

Better understanding public perceptions to pollen seasons provides valuable public health insights. Allergic diseases affect between 10–30% of the global population, with increasing prevalence (Mims, 2014), and its effects are further complicated by climate variability (D’Amato et al., 2020; Reid & Gamble, 2009). Allergies pose a significant economic burden with an estimated $11.2 billion spent on treating allergic rhinitis in the United States in 2005 (Blaiss, 2010), and substantial indirect costs due to decreased productivity, and hidden costs due to comorbidities (Schoenwetter et al., 2004; Blaiss, 2010). On a macro level, public consensus on climate-driven changes in pollen seasons can support governmental and agency adaptation actions. On a micro level, individuals understanding how rising temperatures affect pollen concentrations can facilitate prevention through behavioral changes (Reid & Gamble, 2009).

In order to study the public perceptions of pollen phenology during climate change, we leverage posts from “Twitter” (now “X”), a widely used social media platform. Social media has been shown to provide powerful big data for public perceptions of climate change (Jang & Hart, 2015). Further, the rich textual information gained from social media can provide in-depth insights into public discourses related to climate change (Kim et al., 2021; W. Zhang et al., 2023; Ghermandi et al., 2023; Anderson, 2017). In particular, social media users post observations of plant phenology on large spatial (e.g., continental) and temporal (e.g., decadal) scales (Breckheimer et al., 2020; Gesualdo et al., 2015; Shin et al., 2021; Silva et al., 2018), enabling us to broadly assess public perceptions.

Addressing the potential bottlenecks in climate change communication using a narrative of climate-driven changes in pollen seasons, we assess the accuracy of detection and patterns of attribution of changes in pollen phenology in social media. We ask and answer the following questions:

1. Do the quantity and timing of social media posts correspond to pollen seasons over time and space?
2. Does the user’s attribution of changes in pollen phenology depend on ideology?
3. What is the role of scientific communication in public discussions on the attribution of changes in pollen phenology?

To address the above questions, we focused on the United States in the past two decades and conducted three sets of analyses:

1. We test if the number of pollen-related posts on Twitter reflects temporal and spatial variations in natural pollen phenology, validated by a high-quality *in-situ* pollen concentration dataset.
2. Through an in-depth analysis of texts, we test the relationship between the likelihood of making attribution statements in Twitter posts and estimated user ideology.
3. We further explore the users involved in the dissemination of attribution statements through retweets, with a focus on the role of mass media and scientific experts.

## 2. Methods and Materials

### 2.1. Retrieving pollen-related discussion on Twitter

We retrieved social media discussions on pollen from the Twitter Decahose dataset, a 10% sample of tweets, hosted at the University of Michigan (U-M). During Dec 2–7, 2022, we queried the Twitter Decahose dataset using the PySpark tool “decahose-filter” (Meyer, 2022/2022) with the keyword “pollen” in the Great Lakes High-Performance Computing environment at UM. Queried data covered the time period of 2012 to 2022, with gaps in 2017 and a few other months that had missing or invalid data files. We used only the keyword of “pollen,” but not others such as “allergy”, to obtain a dataset with a clear focus on the perception of changes in the biological system. Queried data contained variables including unique tweet id, date (in the local time zone), user name, user description, user location, language, and text. In order to focus on the Twitter discussion on pollen, we applied the following filters. We removed all tweets with duplicated unique id. We kept only tweets posted in English. We removed tweets that only contain “pollen” in a tag, in a URL, or as a part of a longer word, in order to keep tweets that contain the exact complete word “pollen.”

We applied reverse geocoding to geolocate Twitter users for two purposes: to filter for tweets that likely originated in the US and to further analyze spatial patterns across states within the US (Velardi et al., 2014). User location, self-described by users, often represents the location of tweeting, although not always. Note that many Twitter users choose not to describe their user location, and not all user location information can be identified to a known place (Dredze et al., 2013; Hecht et al., 2011). For users with location information, we took three steps in reverse geocoding. First, we located users to the US by searching for US-related keywords (“us,” “u.s.,” “u.s,” “u.s.a.,” “u.s.a,” “usa,” “united states,” “united states of america,” and “america”) in user location, ignoring case differences. Second, we attempted to locate users to states, based on the proximity of their coordinates to a known US postal code or direct mentioning of the name of a US place (county, city, or state) in a geoname database. Last, we attempted to locate users to large cities (with over 100,000 population) within the US, based on the proximity of their coordinates to a known large city or direct mention of the name of a large city. For the last two steps, we used a python package “twitter-user-geoencoder” (Bian, 2015/2023). Geonames and postal codes used were retrieved from https://download.geonames.org/ on Dec 23, 2022. If a user could be located in any of the three steps, we considered them within the US. We located users to states within the US whenever possible with the last two steps. We acknowledge that this method could only account for possible location at the time of data collection, but not changes over the study period. We kept only tweets posted by a user within the US identified from reverse geocoding. Given that only 66% of users in our dataset provided identifiable locations, our dataset was a subset of all pollen discussions on Twitter in the US, and numerical results should be interpreted in relative terms.

We used AI methodologies to identify distinct and meaningful topics in our pollen-related tweet corpus. We performed topic clustering using the Bidirectional Encoder Representations from Transformers (BERT) topic model (BERTopics), a cutting-edge natural language processing methodology. After creating dense embeddings of tweets based on their semantic meaning using BERT, we used dimensionality reduction using Uniform Manifold Approximation and Projection (UMAP) and clustering through Hierarchical Density-Based Spatial Clustering of Applications with Noise (HDBSCAN). The minimum number of tweets needed to form a cluster was set to 1,152 (1% of the total tweets).

### 2.2. Validating pollen phenology detected by Twitter users

To obtain the temporal trend of pollen phenology detected by Twitter users, we calculated the number of pollen-related tweets (referred to as tweet count) in our dataset on each day. Whenever analyzing tweet count, we adjusted for the expansion in US Twitter use base over time (eqn 1, Fig. S3), making values in the early years comparable to the current ones from approximately 68 million US Twitter users.

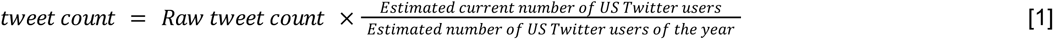

To validate if trends in pollen discussion on Twitter reflect natural pollen phenology, we obtained consistent and accurate pollen concentration data dating back to 2012 from 98 stations in the continental US associated with the American Academy of Allergy, Asthma & Immunology (AAAAI) and the National Allergy Bureau (NAB) (AAAAI, 2022). At stations located throughout the continental US, airborne pollen was sampled nearly daily using volumetric impactor samplers, including either Rotorod or Burkard samplers (Frenz, 1999) and then classified to the genus or family level and counted by NAB-certified operators. The NAB dataset serves as one of the most accurate and consistent data sources for the description and prediction of pollen phenology in public health and ecological research in North America. We cleaned the pollen concentration data to obtain pollen concentration (grains m^-3^) summed over all plant taxa for each day and station.

We compared Twitter pollen phenology characterized by daily tweet count and natural pollen phenology characterized with daily pollen concentration. We assessed the correlation between Twitter and natural pollen phenology on the continental US level. We performed the following data processing steps.

1. We removed combinations of station and year with a total pollen concentration lower than 100 grains m^-3^ to remove possibly erroneous data.
2. We standardized each time series, either US-level tweet count or station-specific pollen concentration, to approximately between zero and one, by dividing all values by the respective 95th percentile. This step allowed comparability of natural pollen phenology across sites and between Twitter and natural pollen phenology.
3. We upscaled natural pollen phenology from the station level to the US level, by averaging the standardized pollen concentration over all stations for each day, removing average values generated by fewer than five data points.
4. We smoothed both US-level Twitter and natural pollen phenology time series with a Whittaker method (Kong et al., 2019), while filling gaps in time series that were no longer than 14 days (Katz et al., 2023).

We then tested the linear correlation between Twitter and natural pollen phenology over all dates.

Spatially, we assessed the correlation between Twitter and natural pollen phenology on the state level. We performed the following data processing steps.

1. We removed combinations of station and year with lower than a total pollen concentration of 100 grains m^-3^ and combinations of state and year with lower than a total tweet count of 100.
2. We upscaled natural pollen phenology from the station level to the state level, by taking the average of pollen concentration over all stations in a state for each day.
3. We estimated long-term Twitter and natural pollen phenology on the state level, using tweet count and pollen concentration on each DOY from 1 to 365 averaged over all years for each state. As pollen concentration data collection had various starting dates, we removed average pollen concentration values generated by fewer than three data points.
4. We smoothed all the time series with the Whittaker method, filling gaps in time series that were no longer than 14 days.
5. We identified for each state the time of spring pollen peak from Twitter pollen phenology as the first DOY that reaches 50% of the cumulative sum (Nilsson & Persson, 1981; Rasmussen, 2002) of the count of pollen-related tweets in the spring pollen window (DOY 32 – 151) (Anderegg et al., 2021). We similarly identified for each state the time of spring pollen peak from natural pollen phenology using the cumulative sum of pollen concentration instead.

We quantitatively tested the linear correlations between phenological metrics (time of spring pollen peak) derived from Twitter and natural pollen phenology. All linear correlations mentioned above were quantified with the Pearson correlation coefficient (*R*) and tested for statistical significance with *t*-tests.

### 2.3. Identifying ideological patterns in attribution

We examined patterns in US Twitter users’ attribution of variations in pollen phenology, specifically to temperature change and climate change. We hypothesized that users’ attributions of changes in pollen season to temperature and climate change do not depend on ideology. Driven by our hypotheses, we defined three groups of Twitter users in the US: a group of users that discuss the phenology of airborne pollen (“pollen group”), a subgroup of users that attributes changes in airborne pollen level to changes in temperature related to temperature (“pollen-temperature group”), and a subgroup of users that attributes changes in airborne pollen level to climate change (“pollen-climate group”). Notably, these three groups were designed to be nested, such that users in the “pollen-temperature group” were by definition in the “pollen group” and that users in the “pollen-climate group” were by definition in the “pollen-temperature group” and “pollen group.” Such as nested design allowed us to easily calculate conditional probabilities of attribution by comparing the number of users between two groups, correcting for baseline distribution of ideology in the whole pollen-related discussion. It also created a gradient of technical complexity in attribution, ranging from a general discussion not necessarily with attribution, to a straightforward attribution to short-term local temperature, and to a convoluted attribution to long-term regional climate.

Every tweet in our sample already contains the keyword “pollen.” We filtered for users in the “pollen-temperature group” by searching for keywords related to temperature change (i.e., “warm”, “warmer”, “warming”, “hot”, and “hotter”); we filtered for users in the “pollen-climate group” by searching for keywords related to climate change (i.e., “climate” and “warming”). We identified all tweets that contained corresponding keywords for the three groups and took a random sample of 1,000 when more tweets were found. As the keyword searching approach was coarse and often did not accurately classify users’ to our defined groups, we read and interpreted the full text in samples from the three groups. We imposed a set of *a priori* criteria to manually label and identify tweets in these three groups. If a user posted at least one of the valid tweets in a group, we identified the user to be in that group. A user might be identified to be in multiple groups, if we found valid tweets posted by them in multiple groups.

We estimated the ideology of identified users in these three groups using an established method of correspondence analysis (Barberá et al., 2015). This method uses commonly followed Twitter accounts, not necessarily political, to project users to a latent ideological space, assuming that users who follow a similar set of accounts are more likely to have similar ideology. A more positive ideology score corresponds to relatively conservative ideology, and a more negative ideology score corresponds to relatively liberal ideology. We retrieved users’ followed accounts using Twitter’s API between Mar 29 and 31, 2023. Our ideology scoring was implemented with functions adapted from the R package “tweetscores” (Barberá et al., 2015; Barberá, 2021). This method leverages the principle of homophily, a well-established and simplified way to predict political ideology. It uses correspondence analysis to project users into a latent space based on accounts they follow. Based on the relative positions of users in the latent space, they are given ideological scores that indicate their relative position on the liberal-conservative spectrum. Note that we could not estimate the ideology of all users, because many have deleted their accounts, did not give permission to access friends list, or did not follow any popular accounts.

We examined the effects of ideology on attribution in two ways. First, we qualitatively compared the probability density distribution of user ideology in three groups, after smoothing histograms with Gaussian kernels of 0.2 bandwidth. Second, we quantitatively tested for the correlation between the conditional probability of attribution and ideology. We calculated both conditional probability (attribution to temperature change given discussion of pollen phenology and attribution to climate change given discussion of pollen phenology) for each bin of ideology (eqn 2). We binned ideology scores from –2 to 2, at intervals of 0.5, leading to eight bins in total.

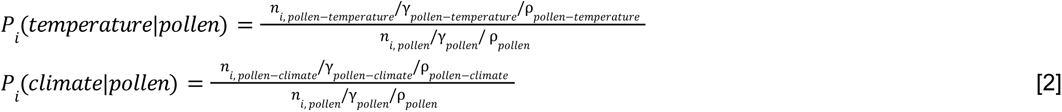

Here, *i* refers to the bin of ideology score, ranging from 1 to 8. *γ* refers to the retrieval rate for a group when we only successfully retrieved ideology for a subset of identified users and *ρ* refers to the sampling rate for a group when we took a random sample for manual labeling (eqn 3).

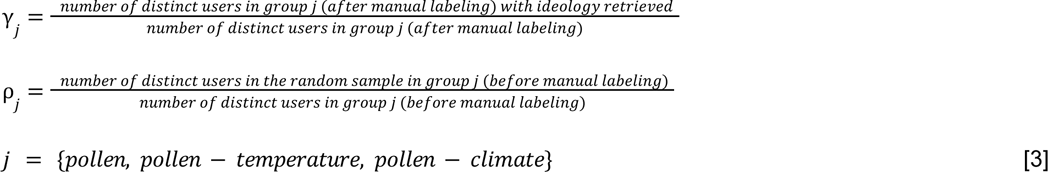

We tested if the conditional probability *P_i_* was linearly correlated to the bin of ideology score. Because of the limited number of users in the sample, we performed bootstrapping to obtain more robust relationships, accounting for uncertainty in conditional probability. Specifically, we resampled the entire estimated user ideology dataset with replacement 1000 times. We then calculated 1000 sets of conditional probability across bins of ideology scores using the bootstrapped samples. We performed linear regression using conditional probability against the bin of ideology score (converted to integers from 1 to 8) for each set of conditional probability and summarized the medians and 95% intervals of slopes of regression (*β*). We used all fitted linear regression models to generate the medians and 95% intervals of predictions.

### 2.4. Visualizing the flow of information

In order to explain possible ideological patterns in the attribution of pollen phenology, we analyzed the flow of information, with a focus on the role of scientific communication. We characterized the flow of information specifically by actions of retweeting, using the user being retweeted as a proxy for the source of information, and the user posting the retweet as a proxy for the destination of information. In order to identify the major types of information flow in the discussions, we read usernames and user descriptions and manually classified the type of user, either the source or destination of information, into four mutually exclusive *a priori* categories.

A) “media”: Accounts that had the main purpose of disseminating information to the public, such as TV stations, weather stations, and newspapers. Individuals who work in the media and use their accounts mainly for information dissemination (instead of for personal opinions) are also included in this category, such as news reporters, meteorologists who work at news stations, and magazine editors.
B) “experts”: Professionals who conduct research in allergy, phenology, meteorology, or climate change. Organizations with a group of experts, such as the American Academy of Allergy, Asthma, and Immunology, are included in this category.
C) “others”: Accounts that are not classified as either media or experts

a. “other organizations”: Accounts that are not classified as either media or experts, and are used collectively by a group, such as a university or a climate activist group.
b. “other individuals”: Accounts that are not classified as either media or experts, and are used by an individual.

We were particularly interested in the roles of scientific communication, so we explicitly represented “media” and “experts” users in our dataset, with all other accounts being classified into the “others” category. The choices of these categories were driven by theories and hypotheses on the possible roles of mass media and scientists in the climate change discourse (Carvalho, 2010; McCright et al., 2013; Hmielowski et al., 2014). As individuals and organizations use Twitter very differently, we distinguished “other organizations” and “other individuals.” We visualized the flow of information from source to destination in the three nested groups of discussions with Sankey diagrams. We performed Pearson’s chi-squared test on the numbers of source or destination users in four categories among three groups of discussion, to compare if the proportions for the four categories differ significantly.

All data analyses were performed in *R* v. 4.2.0 (R Core Team, 2023), except that topic modeling was performed in *python* v. 3.10.12 (Python Software Foundation, 2024).

## 3. Results

### 3.1. Twitter users accurately detect pollen phenology

We compiled a dataset of 190,473 unique English-language tweets containing the keyword “pollen” posted in the continental US from 2012 to 2022 (Fig. S1), with 177,722 that could be geographically resolved to states. This dataset contained tweets on relevant and diverse topics including pollen allergies, observations of nature, weather forecasts, and car cleaning (Fig. S2). Public discourse on pollen on social media is highly relevant to pollen phenology, but is also diverse and nuanced. We found strong seasonality in the mention of pollen on Twitter in the US (Fig. 1a, Figs. S3, S4, S5), with the daily tweet count sharply increasing in March, peaking in early April, and gradually decreasing until late June. In our dataset, 64% of the tweets were posted in the spring pollen window from day-of-year (DOY) 32 (Feb 1) to DOY 151 (May 31) (Anderegg et al., 2021).

**Fig. 1:**
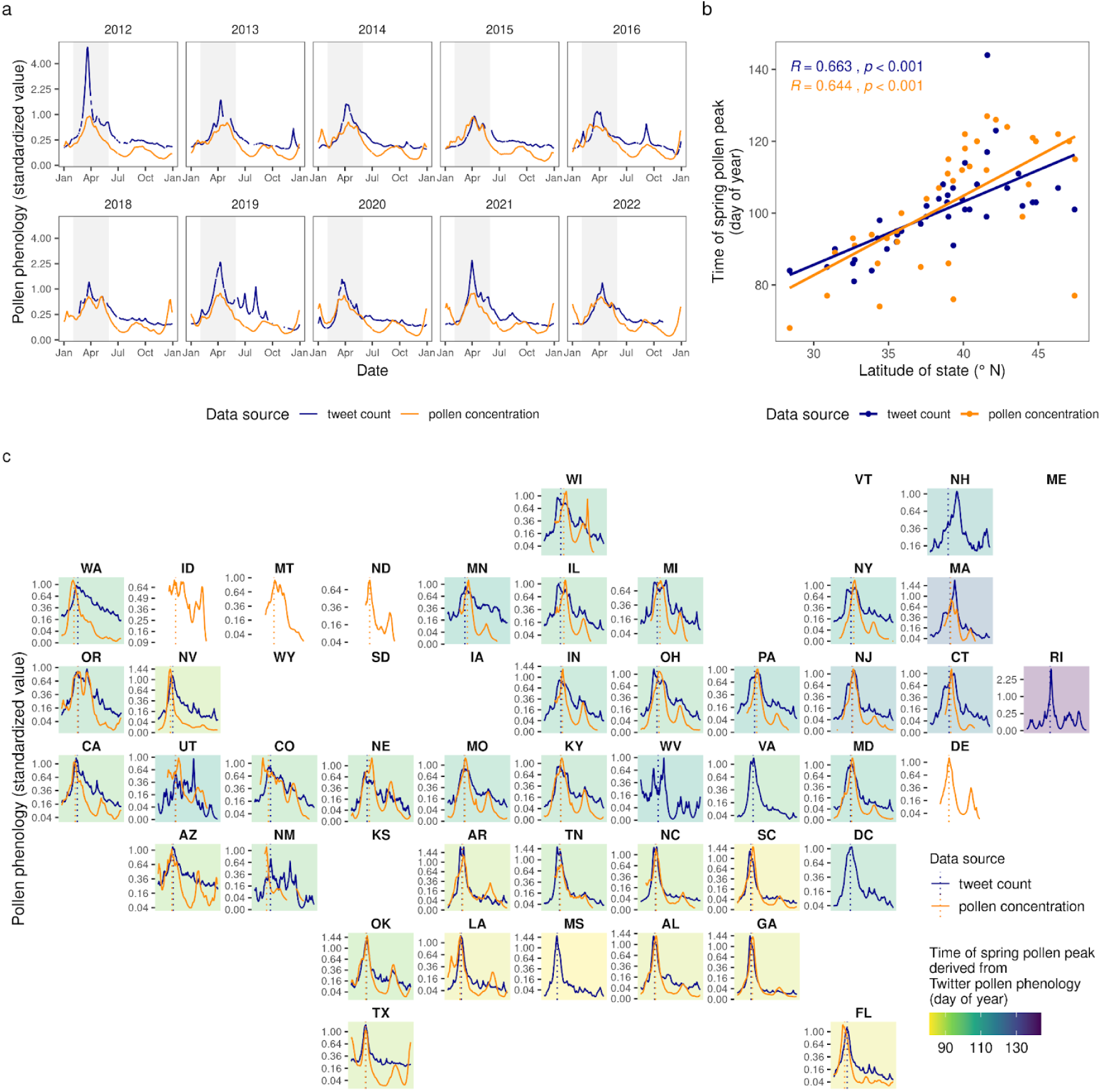
Twitter pollen phenology aligns with natural pollen phenology over time and space. **a**, Trends in Twitter pollen phenology and natural pollen phenology in the continental US from 2012 to 2022 (missing data in 2017), derived from the count of pollen-related tweets (in our compiled dataset) and pollen concentration (from pollen counting stations associated with the National Allergy Bureau, NAB), respectively. Gray shades delineate a defined window for spring pollen peaks, from day-of-year 32 to 151 (February to May). **b**, Both Twitter and natural pollen phenology exhibit latitudinal gradients. Trend lines show the correlation between the latitude of the geometric centroid of a state and the time of spring pollen peak (day of year) derived from Twitter and natural pollen phenology, respectively. Pearson correlation coefficient (*R*) and *p*-value (*t*-test) are shown. **c**, Spatial variation of pollen phenology across states in the US. Lines show long-term average Twitter and natural pollen phenology in each state. Dotted lines and colors of panels show the time of spring pollen peak derived from Twitter and natural pollen phenology (day of year). A brighter color of the panel indicates an earlier spring pollen peak derived from Twitter pollen phenology in the state.

We validated pollen phenology data derived from Twitter with a long-term continental-scale standard dataset collected from pollen counting stations associated with the National Allergy Bureau (NAB) (AAAAI, 2022) (Figs. S6, S7, S8). Comparing pollen phenology characterized by the count of pollen-related tweets (referred to as “Twitter pollen phenology”) and pollen phenology characterized by pollen concentration (referred to as “natural pollen phenology”), we examined how accurately pollen discussions on Twitter reflect temporal and spatial variations in pollen phenology. There were similar temporal trends between Twitter and natural pollen phenology between 2012 and 2022 for the US (Fig. 1a). The most dominant pollen peaks in spring were evident in comparing accounts of Twitter and natural pollen phenology, while the minor Fall and New Year peaks were not captured on Twitter. There was a significant correlation between pollen phenology derived from the two sources (*p* < 0.001, *t*-test, Fig. S9). Both Twitter and natural phenology exhibit a latitudinal gradient over space, with southern states experiencing earlier pollen seasons compared to northern states (Fig. 1b, 1c, Figs. S10, S11). Across 29 states with data available from both Twitter and NAB datasets, we found a significant correlation between the time of spring pollen peak derived from Twitter and natural phenology (*p* < 0.001, Fig. S12). These results together supported that the discussion of pollen on social media reflects natural pollen phenology, both over time and across space.

### 3.2. Attribution to climate change depends on political ideology

We tested the hypotheses that users attributing the changing pollen season to temperature change or climate change is not correlated with ideology. We defined three hypothesis-driven nested groups of users based on how their tweets attribute changes in pollen phenology: those that discuss pollen phenology in general (e.g., “this pollen is killing me”), those that attribute changes in pollen phenology to temperature change (e.g., “pollen level is high and warm weather will make it worse”), and those that attribute them to climate change (e.g., “global warming could make your pollen allergies a lot worse”) (Fig. 2a). We followed an *a priori* coding scheme to manually label random samples of tweets with selected keywords, in order to accurately identify tweets that fit the criteria of the three groups, screening out tweets that share similar keywords but had different meanings (Note S1). We were able to assemble a dataset of 829 users’ ideology scores. We then estimated the ideology of each user in the three groups by retrieving the accounts they follow and performing a correspondence analysis that mapped users to a latent ideology space (Barberá et al., 2015).

**Fig. 2:**
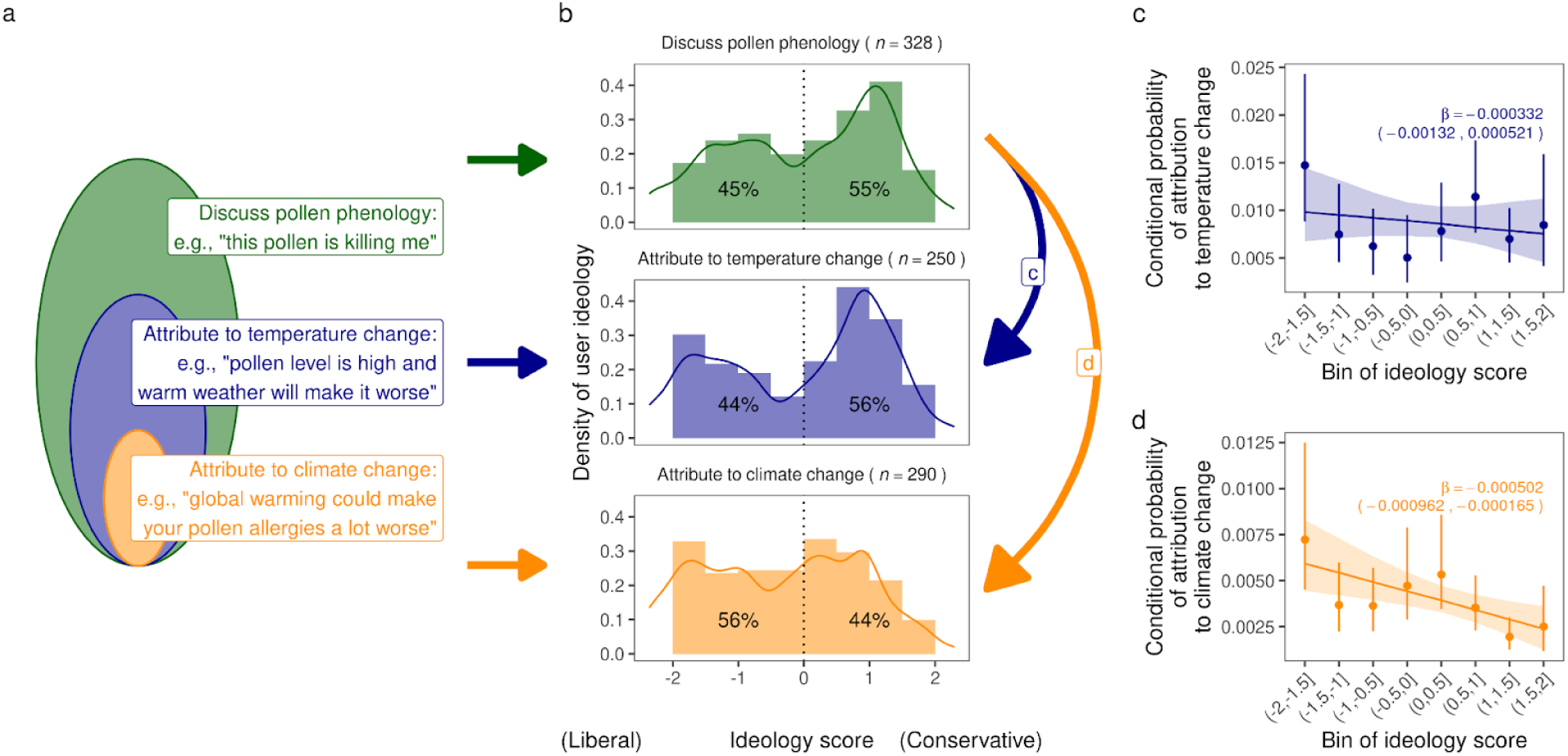
Twitter users’ attribution of pollen phenology was likely ideologically structured. **a**, Venn diagram showing the relationship between three groups of continental US Twitter users defined for this study. Hypothetical tweets that reflect the definitions of three groups are shown, as original texts could not be shown for privacy concerns. **b**, Density (histogram) and kernel density estimates (smoothed histogram) of ideology score in three groups of users in our compiled dataset. Number of users with estimated ideology score (*n*) in each group and percentages of users with negative and positive ideology scores (relatively liberal and conservative, respectively) are shown. **c&d**, Correlation between the conditional probability of a user attributing changes in pollen phenology to a specific driver (temperature change or climate change) given general interest in pollen phenology and the bin of ideology the user was in. Medians and error bars of estimated conditional probability were obtained from bootstrapping user ideology scores 1000 times. Lines and ribbons show predicted linear trends and 95% intervals using linear models fitted with bootstrapped samples. Medians and 95% intervals of the slopes (*β*) are shown.

We showed a qualitative difference in users’ ideology among the three groups that attribute changes in pollen phenology differently (Fig. 2b). The probability density distribution of user ideology in all three groups showed bimodal distributions, with two peaks corresponding to relatively conservative (positive ideology score) and relatively liberal users (negative ideology score). In the group that discusses pollen phenology and the group that makes attribution to temperature change, there were more conservative than liberal users. The ideological distributions in these two groups were consistent with the fact that a large proportion of tweets in our dataset came from users in conservative states such as Texas (13%) and Georgia (10%) with higher airborne pollen concentrations. In contrast, in the group that makes an attribution to climate change, there were more liberal than conservative users.

We quantitatively compared how the attribution of changes in pollen phenology depend on political ideology. Enabled by the unique nested design of three focal groups (Fig. 1a), we used the ratio of number of users between groups (Fig. 2b) to estimate the conditional probabilities of attribution given general interest in pollen phenology (Fig. 2c&d). The calculation of conditional probability corrected for the unequal number of liberal and conservative users in the whole pollen-related discussion. A significant correlation between conditional probabilities of attribution and ideology would suggest ideologically-structured attribution. Our results show that the conditional probability of attributing changes in pollen phenology to climate change given general interest in pollen phenology was significantly correlated to the bin of ideology a user was in (*p* < 0.05), with more liberal users being more likely to make the attribution to climate change (Fig. 2d). In contrast, we did not find significant correlation between attribution to temperature change and user ideology (Fig. 2c). The shifting distribution of ideology and ideology-dependent conditional probability suggested that social media users’ attribution of changes in pollen phenology to climate change are structured depending on political ideology.

### 3.3. Scientific communication dominates discussions on climate change impacts

We sought to infer processes behind the observed ideological pattern by examining the flow of information in the three nested groups of discussions (Fig. 2a), with the hypothesized potential roles of mass media and scientists. We focused on retweets because they revealed the source (user being retweeted) and destination (user posting retweet) of information. In both sources and destinations, we examined the presence of users that represent media, experts (on allergy, phenology, meteorology, or climate change), other organizations (not classified as either media or experts and are used collectively by a group), and other individuals (not classified as either media or experts and are used collectively by an individual). We were able to classify users involved in a total of 524 retweets. This allowed us to compare the roles of scientific communication initiated by media or experts and public engagement initiated by non-media and non-experts.

We revealed gradual changes in the dominant users across the three groups of discussions, moving from a general discussion of pollen phenology not necessarily involving attribution to a technically convoluted attribution of changing pollen seasons to climate change. The general discussions on pollen were dominated by communication among individuals, such as retweeting complaints about the allergy season (Fig. 3a). Individuals often retweet posts by the media, such as pollen level forecasts. In the more specific discussions related to temperature changes, we still found a large proportion of communication among individuals (Fig. 3b). However, there was a greater importance of media, followed by experts, in spreading information on the correlation between changing pollen concentration and temperature to individuals. In the discussion related to climate change, the roles of media and experts were even more dominant, being 46% and 20% of the sources of information on the impacts of climate change on changing pollen season, respectively (Fig. 3c). Although individuals were the dominant destinations of information across all groups, it is notable that in the two groups that make attribution to specific drivers, there was considerable communication among media and experts. For example, more than half of the posts by experts on climate-driven changes in pollen seasons were retweeted by fellow experts (e.g., American Academy of Allergy, Asthma, and Immunology (AAAAI) and the director of the USA National Phenology Network, Dr. Theresa Crimmins).

**Fig. 3.**
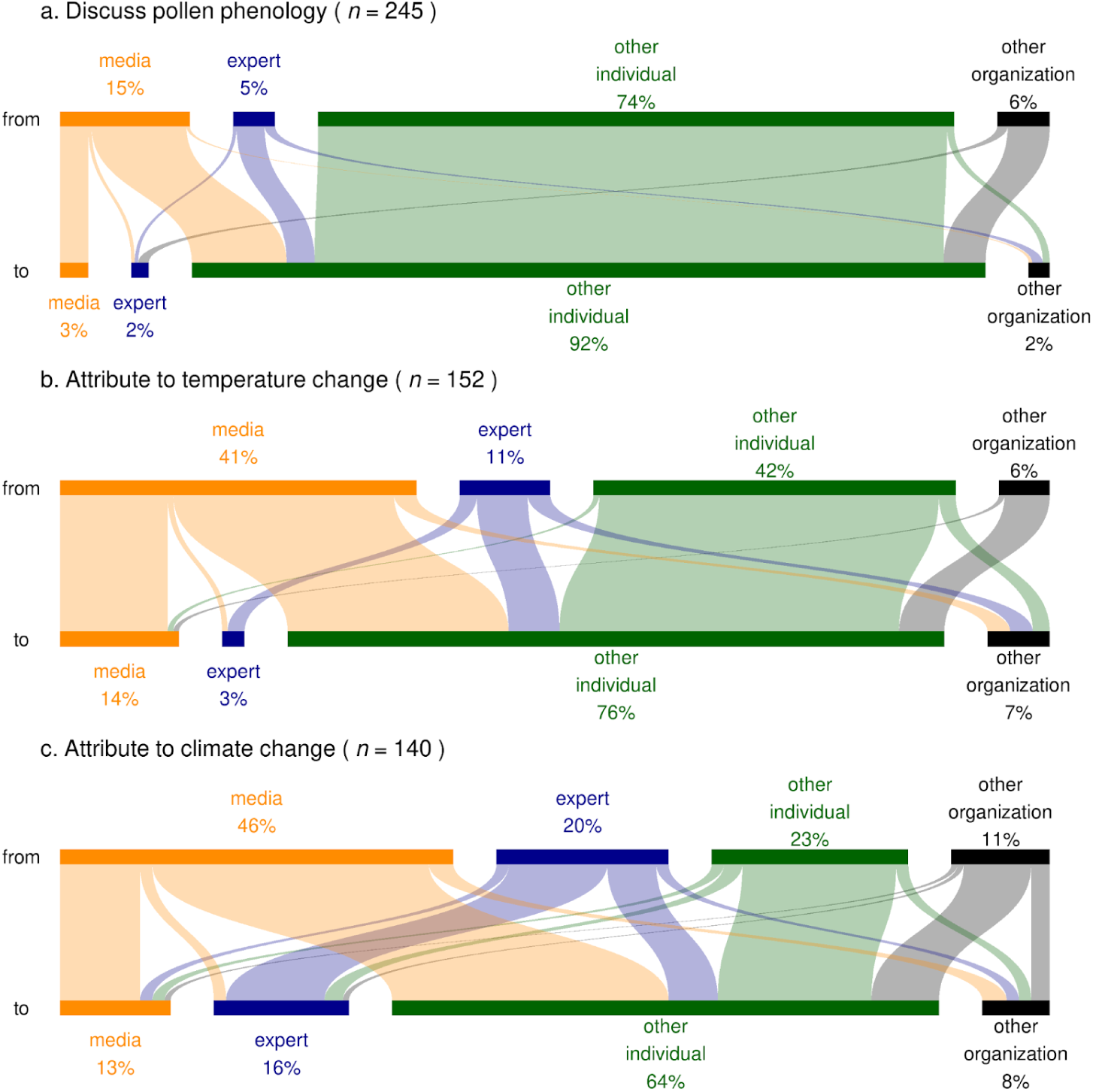
Role of scientific communication on Twitter users’ discussions on pollen phenology. The series of Sankey diagrams show the flow of messages from the source (users being retweeted) to the destination (users posting retweets) in our compiled dataset. We manually classified users into four types: media, experts (on allergy, phenology, meteorology, or climate change), other individuals, and other organizations. The number of retweets (*n*) in each group is shown. Compared to (**a**) the discussions on pollen phenology and (**b**) the attribution of changes in pollen phenology to temperature change, (**c**) the attribution of changes in pollen phenology to climate change were more dominated by scientific communication from media and experts, compared to public engagement from non-media and non-experts. Proportions of the four categories of users, both as source and destination, were significantly different among the three groups of discussion (Pearson’s chi-squared test, *p* < 0.05).

## 4. Discussion

This study focuses on the human dimension of climate change by examining social media users’ perceptions of changing pollen phenology. Pollen discussions on Twitter in the US showed strong seasonality that aligned with natural pollen phenology over time and space, suggesting that social media users are likely to accurately detect variations in pollen phenology. Analyzing a subset of tweets and users, we revealed an ideological pattern in attributing variations in pollen phenology to climate change, with liberal users more likely to agree that climate change affects pollen phenology. As the technical complexity of the attribution increases, we found an increasingly dominant role of scientific communication initiated by the media and experts. Our study contributes to our understanding of human perceptions of one of the most visible climate change impacts, with implications for message framing for climate change communication and public health interventions.

Our findings validate on large spatiotemporal scales that social media users respond timely and predictably to variations in pollen phenology, including rises and falls of pollen concentration in the US, and the sequence of pollen season across US states. Uniquely, we revealed consistency in trends not only over time but over space despite a limited sample that could be resolved geographically (Cowie et al., 2018), suggesting that changes in social media posts were not only caused by spatially uniform factors, such as national media coverage. Our results were supported by previous findings, for example, that Twitter discussions on allergy symptoms and antihistamine drugs aligned with airborne pollen concentrations in 2017 at the US and state levels (Gesualdo et al., 2015). Beyond social media, the close connection between human perceptions and pollen phenology is visible from internet searches (Hall et al., 2020), medication sales (Sheffield et al., 2011), and hospital visits (Lappe et al., 2023; Wakamiya et al., 2019). Pollen phenology is an exceptional representation of broader changes in biological systems that are captured by the public, other examples being vegetation greenness (Silva et al., 2018) and fall phenology (Shin et al., 2021). Changes in biological systems might be more accurately perceived by the public compared to changes in the physical systems, including weather, weather anomalies, and extreme weather events, which have been shown to be highly influenced by expectations, memory limitations, and cognitive biases (Hamilton & Stampone, 2013; Howe & Leiserowitz, 2013; Kirilenko et al., 2015; Moore et al., 2019). The accurate detection of pollen phenology by Twitter users sets a foundation for communicating climate change impacts to the public through experiences of changes in pollen season.

Our study revealed both challenges and opportunities for leveraging pollen phenology as a message to communicate the personal impacts of climate change. While the public is sensitive to changes in pollen seasons, not all have connected these personal experiences with climate change. Indeed, our findings suggested that the attribution of pollen season changes to climate change depended on ideology, consistent with the ideological bias in other climate change perceptions (Cacciatore et al., 2016; Dunlap et al., 2016; Falkenberg et al., 2022; Jang & Hart, 2015; McCright et al., 2016; McCright & Dunlap, 2011; Weber, 2010; Yeo et al., 2017). Previous research has suggested various factors that contribute to such ideological bias, including the influence of media and opinion leaders (Brulle et al., 2012; Carmichael et al., 2017), the politicization of climate-related issues (Bolsen & Druckman, 2018), underlying ideological worldviews (Kahan et al., 2012), trust in scientific institutions (Hmielowski et al., 2014), and the role of social networks (Hart et al., 2015; Williams et al., 2015). These factors often interact and reinforce each other, perpetuating biased perceptions. We acknowledge that our correlational analysis does not exclude the effects of a range of factors that are correlated with ideology, such as income, education, and religiosity (Ballew et al., 2020). In our study, we observed the influence of media and experts, in shaping the discourse on climate change impacts on pollen phenology. Biases in the media and opinion leaders (Boykoff & Boykoff, 2004; Cacciatore et al., 2016), reflected in the lack of representation of extreme right-wing users (Fig. 2a), may amplify political echo chambers in this social media discourse on climate-driven changes in pollen seasons (Anderson, 2017; B. Lee et al., 2023).

We also identified encouraging signs that discussions on pollen phenology have the potential to reduce the ideological divide on climate change perceptions. The public discourse on pollen is highly diverse and nuanced, including the impacts of pollen on individuals’ daily lives beyond allergies, as seen from the complaints about car washing after pollen deposition (Fig. S2). Interpersonal discussion of pollen season on social media might encourage the learning and acceptance of relevant knowledge about climate change (Goldberg et al., 2019). We observed a high level of interest among relatively conservative users in pollen discussions, which was consistent with the high pollen concentration in conservative states (Manangan et al., 2021). The association between pollen and temperature resonated with users across different ideologies, offering an opportunity to establish common ground in climate change communication (Bloodhart et al., 2015; Shao, 2016). The topic of climate-driven changes in pollen seasons, being discussed in a bipartisan network without strong salience of partisanship, has the potential to eliminate belief polarization (Guilbeault et al., 2018). The influential role of scientific communication initiated by media and experts offers opportunities for scientists to contribute to the discussion of this technically complex topic by increasing media exposure and public outreach. Despite the foreseeable challenges (Egan & Mullin, 2012), messages revolving around pollen phenology hold promise in strategically bridging the ideological divide in public perceptions about climate change.

Beyond climate change communication, there are implications for public health intervention design. The spatiotemporal alignment between perceived and natural pollen seasons suggests that changes in pollen season are likely experienced on individual user levels, motivating communication of pollen-related personal health risks. The link between higher temperature and pollen concentration was similarly received across ideology, further supporting the potential effectiveness of science-driven precision health interventions such as reducing outdoor activities on hot days. Nevertheless, the ideological difference in perceiving the link between pollen seasons and climate change asks for tailored public health messages to foster long-term behavioral and policy shifts.

We recognize a few limitations in our study. To obtain a focused and consistent dataset, we limited our scope by choices such as the keyword (“pollen”), geographical area (United States), language (English), and data source (Twitter). Future research using allergy-related keywords (Gesualdo et al., 2015), global extent, multiple languages, and diverse data collection methods will help test the generalizability of our findings. Due to the large number of pollen-related tweets, we could not discern tweets generated by individuals from the public and those generated by groups, except for a subset of tweets in the last analysis (Pickering & Norman, 2020). Distinguishing between these posts would have enabled us to understand how the seasonality of pollen-related discussion by individuals is influenced by exposure to media (Egan & Mullin, 2017), which may exhibit its own seasonality in news coverage. To ensure data quality, we manually screened random tweets and users, leading to limited sample sizes for the analyses of political ideology and scientific communication. Nevertheless, our sample sizes were above the median sample size of similar studies (Cruz, 2017). Due to the limited contextual information provided by Twitter data, as compared to resources like surveys or dialogue-rich social platforms, we were not able to disentangle the attribution influenced by ideas propagated by the media at the time from that solely stemming from users’ inherent long-term understanding (Carvalho, 2010). The study also faced limitations imposed by the Twitter policies that protected the privacy of users, challenging our ability to comprehensively measure user location and identity (Pickering & Norman, 2020). This limited the spatiotemporal resolution of the detection accuracy analysis and restricted the sample size for the attribution pattern analyses. We anticipate future research on the perceptions of biological systems using social media data from other platforms.

## 5. Conclusion

This study represents a step towards achieving a more comprehensive understanding of the intricate relationship between climate change, pollen phenology, and public perceptions. Messages revolving around climate-driven changing pollen phenology are likely to help overcome a key bottleneck in communicating climate change impacts: unlike the inaccurate accounts of weather, the public is sensitive to changes in pollen phenology. Nevertheless, such messages still face the other bottleneck: the way changes in pollen phenology are attributed to climate change is shaped by ideology, similar to the polarization of public perceptions on many other climate change topics. Scientific communication by media and experts dominates this ideologically-structured attribution, pointing to both possible mechanisms and potential paths forward.

## Supplemental information

Figures S1–S12 and Note S1.

## Acknowledgments

We thank Chuangbo Tong, Luke Hamilton, and Cait Schilt for exploratory data analyses. We thank members of the University of Michigan Institute for Global Change Biology and Zhu Lab for their constructive comments. This work was supported by the Eric and Wendy Schmidt AI in Science Postdoctoral Fellowship, a Schmidt Sciences program (YS, NF), and the National Science Foundation [grant numbers 2045309 (CAREER)] (KZ, YS).

## CRediT authorship contribution statement

**Yiluan Song:** Conceptualization, Methodology, Data Curation, Formal Analysis, Validation, Visualization, Writing – Original Draft, Writing – Review & Editing

**Adam Millard-Ball:** Conceptualization, Writing – Review & Editing

**Nathan Fox:** Formal Analysis, Writing – Review & Editing

**Derek Van Berkel:** Writing – Review & Editing

**Arun Agrawal:** Writing – Review & Editing

**Kai Zhu:** Resources, Conceptualization, Writing – Review & Editing

## Data and code availability

We retrieved Twitter discussions on pollen from the Twitter Decahose dataset hosted at the University of Michigan. Twitter data were queried during Dec 2–7, 2022. We ceased all querying of Twitter data after the end of U-M’s license for Twitter Decahose dataset and Twitter’s transformation to X. For legal and ethical considerations, raw data (tweet text and user information) will not be released. We obtained pollen concentration data from pollen counting stations associated with the National Allergy Bureau. Pollen concentration data were received on Apr 25, 2023. Processed data to reproduce all analyses will be available upon request.

