## Supplementary Information for "Biological changes, political ideology, and scientific communication shape human perceptions of pollen seasons"

*Kai Zhu

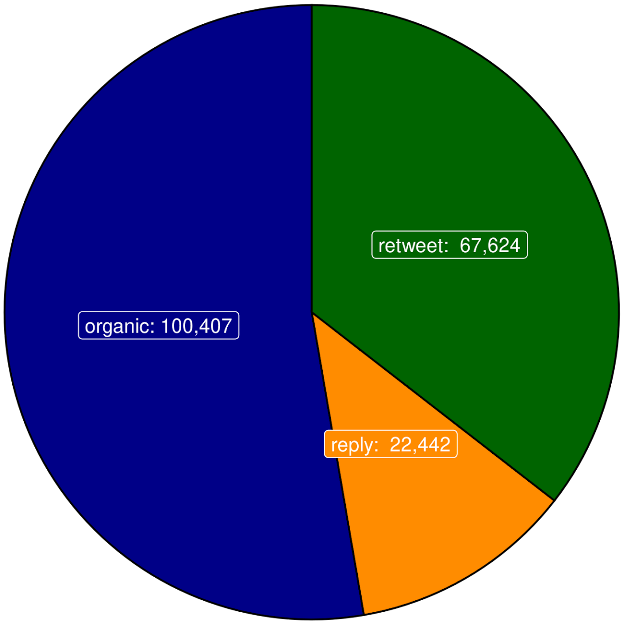


**Fig. S1.** Tweets that qualified searching criteria and were included in our dataset were from three categories (organic, reply, or retweet). The number of tweets is shown in each category. Our dataset was a subset of all pollen discussions on Twitter within the US.

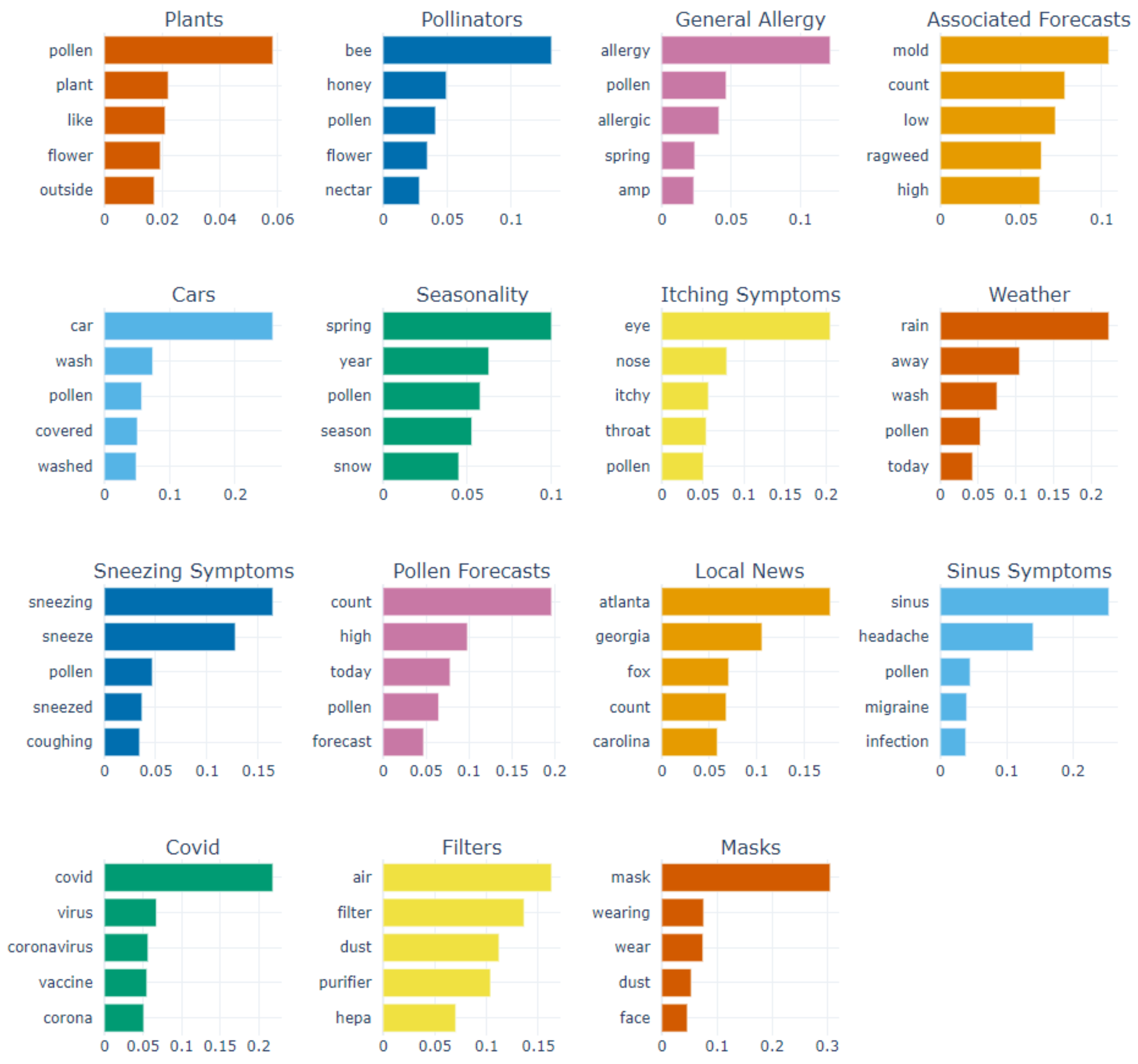


**Fig. S2.** We identified 15 distinct topic clusters in pollen-related tweets. For each topic cluster, we show the five most important topic words and their scores. We named each topic cluster according to these topic words. The three most prominent clusters were “Plants,” “Pollinators,” and “General Allergies,” containing 17,501, 13,153, and 13,061 tweets respectively. These clusters provided insightful categorization, with “Plants” predominantly encompassing tweets about the plants and flowers, “Pollinators” focusing on discussions around bees and other pollinators, and “General Allergies” capturing tweets related to general allergic reactions and health concerns. A portion of the dataset, comprising 32,883 Tweets, did not align distinctly with any of the identified clusters. This subset of tweets may represent a more diverse range of topics or nuanced discussions that did not converge into clear thematic groups based on the applied clustering parameters.

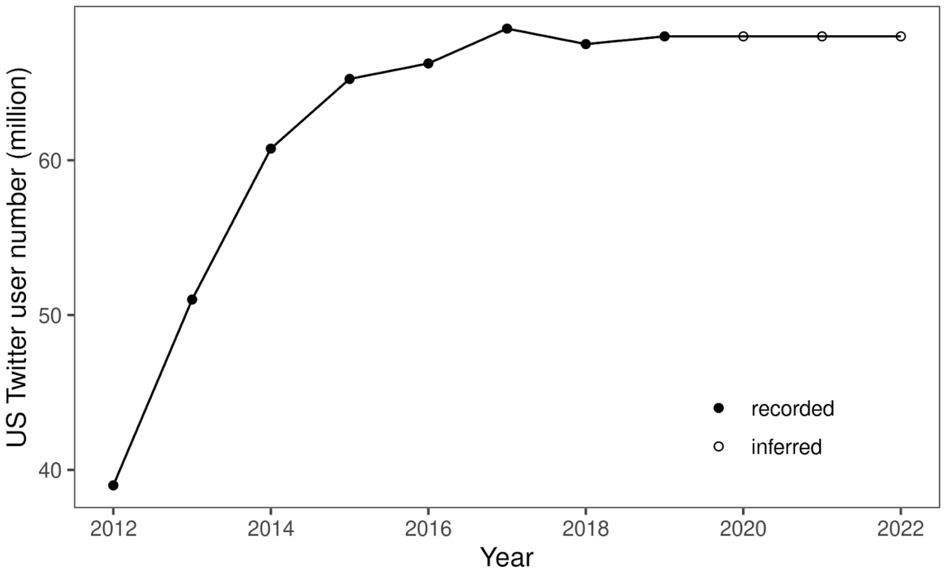


**Fig. S3.** We adjusted the number of pollen-related tweets for Twitter's user growth over the years. We estimated the number of active Twitter users in the United States (US) from 2012 to 2022 using a dataset of the number of monthly active Twitter users in the US from 1st quarter of 2010 to 1st quarter of 2019 on Statista (https://www.statista.com/). We assumed the same number of users after 2019.

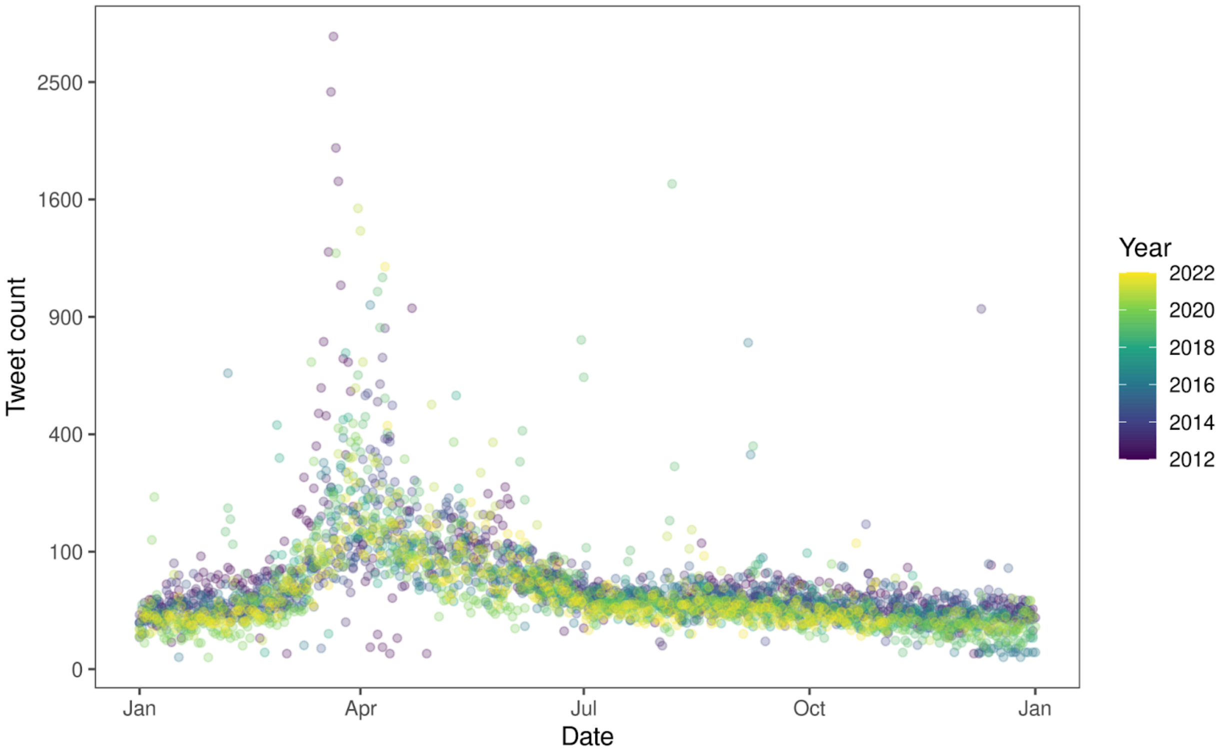


**Fig. S4.** The count of pollen-related tweets in our compiled dataset on each day from 2012 to 2022. The colors indicate different years. Our compiled dataset was a subset of all pollen discussions on Twitter within the US.

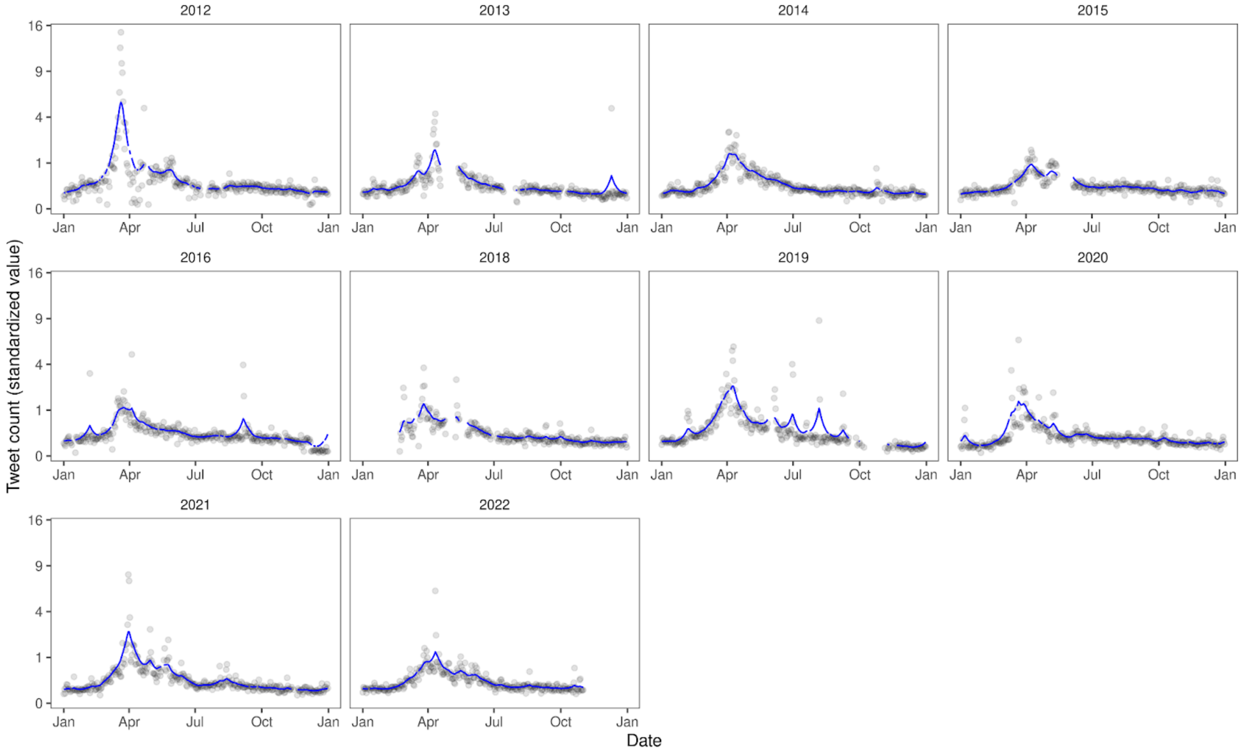


**Fig. S5.** Standardized count of pollen-related tweets in our compiled dataset on each day from 2012 to 2022, characterizing Twitter pollen phenology. Black points and blue lines indicate values before and after smoothing and gap-filling, respectively. Our dataset was a subset of all pollen discussions on Twitter within the US.

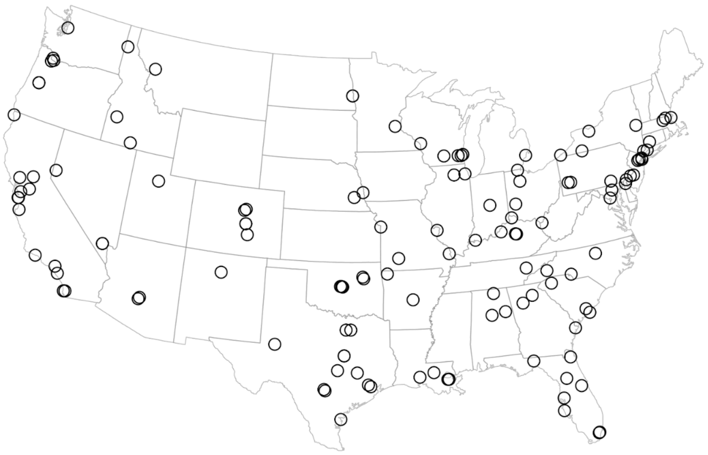

**Fig. S6.** Map of pollen counting stations associated with the National Allergy Bureau (NAB) within the continental United States that contributed to pollen concentration data used in this study.

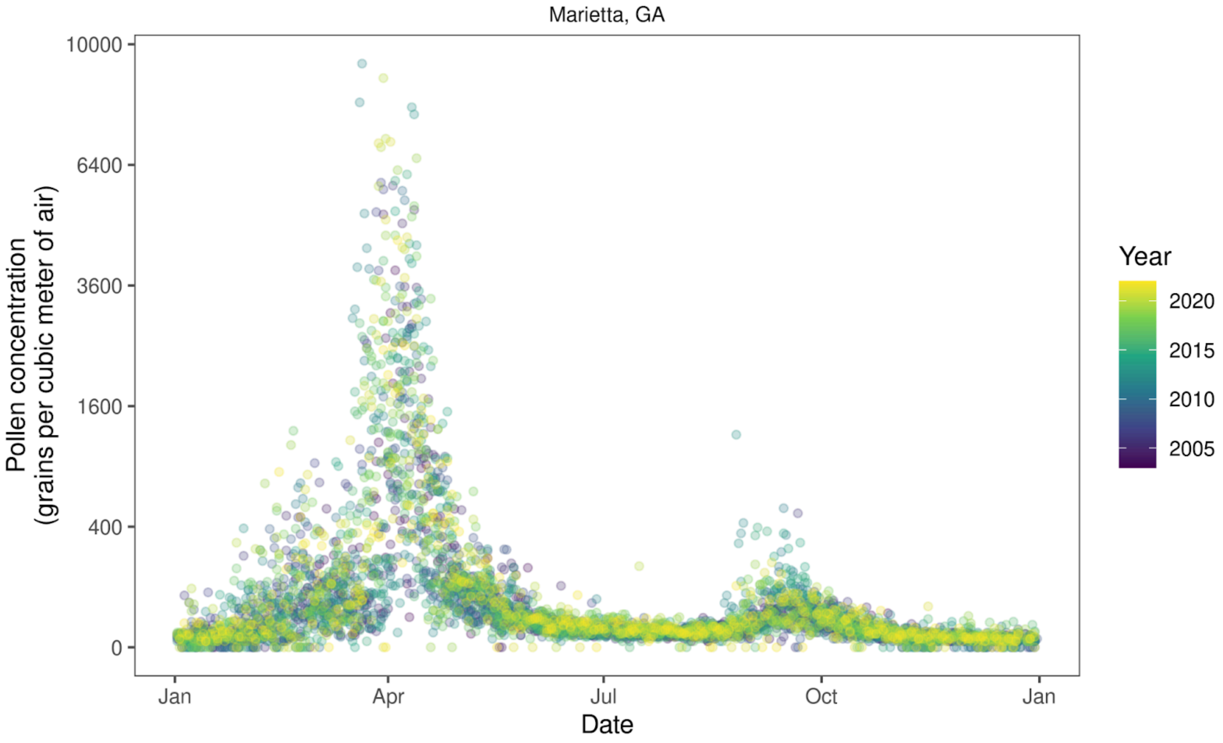


**Fig. S7.** Pollen concentration (grains per cubic meter of air) on each day. Data from one pollen counting station in Marietta, Georgia (GA) associated with the National Allergy Bureau (NAB) are shown here as an example. Colors indicate different years.

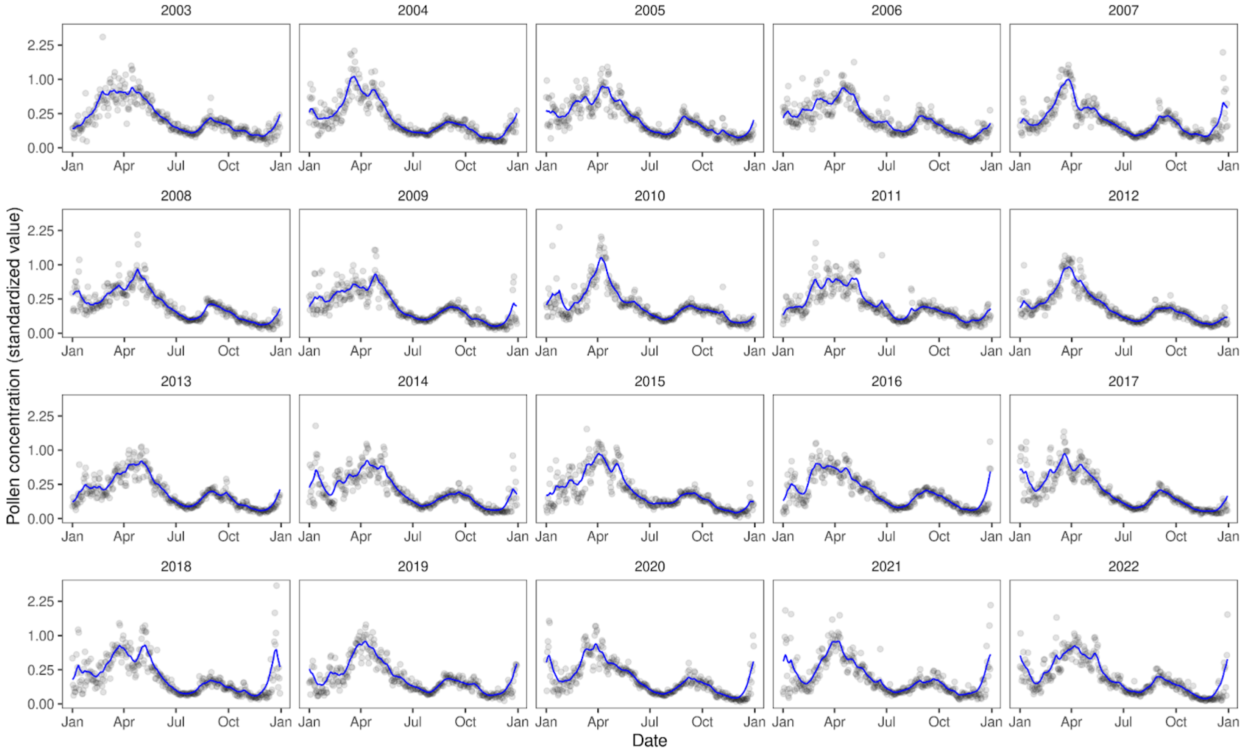


**Fig. S8.** Standardized pollen concentration averaged to the US level on each day from 2003 to 2022, characterizes natural pollen phenology. Raw pollen concentration data were from pollen counting stations associated with the National Allergy Bureau (NAB) within the continental United States. Black points and blue lines indicate values before and after smoothing and gap-filling, respectively.

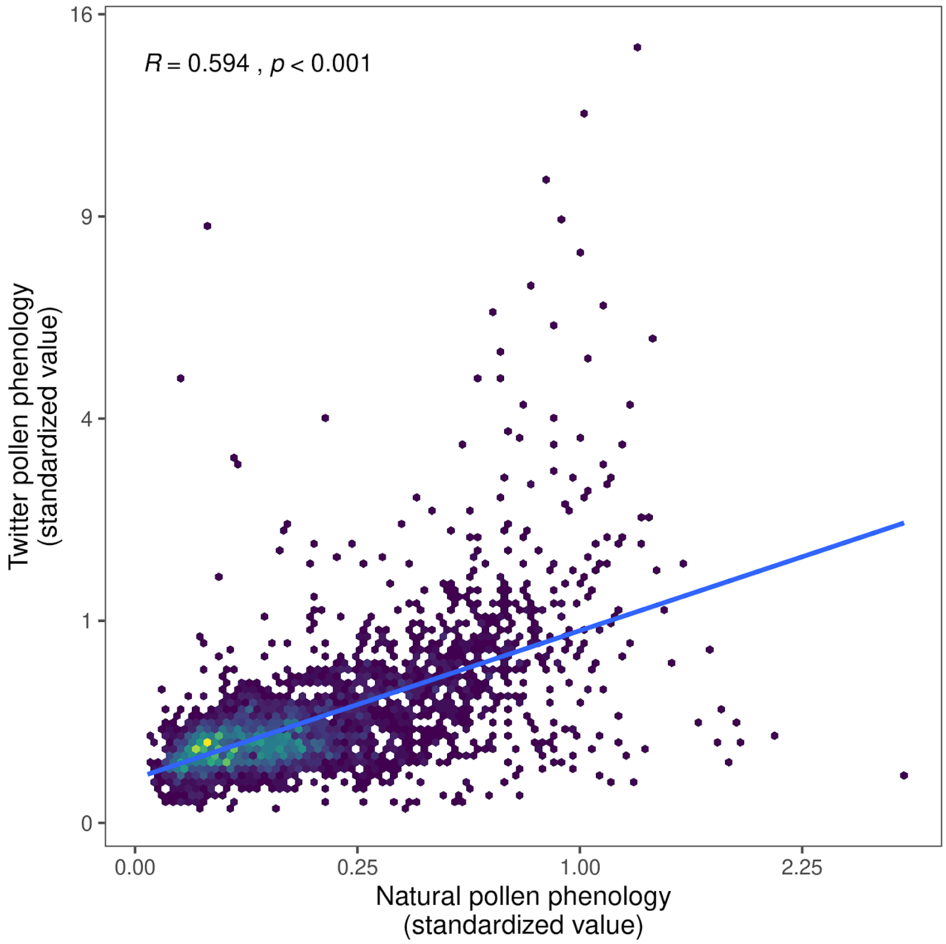


**Fig. S9.** Heat plot showing the correlation between natural pollen phenology derived from pollen concentration and Twitter pollen phenology derived from count of pollen-related tweets. A brighter color indicates a greater number of data points. Trend lines, Pearson correlation coefficient ($R$), and $p$-value ($t$-test) are shown.

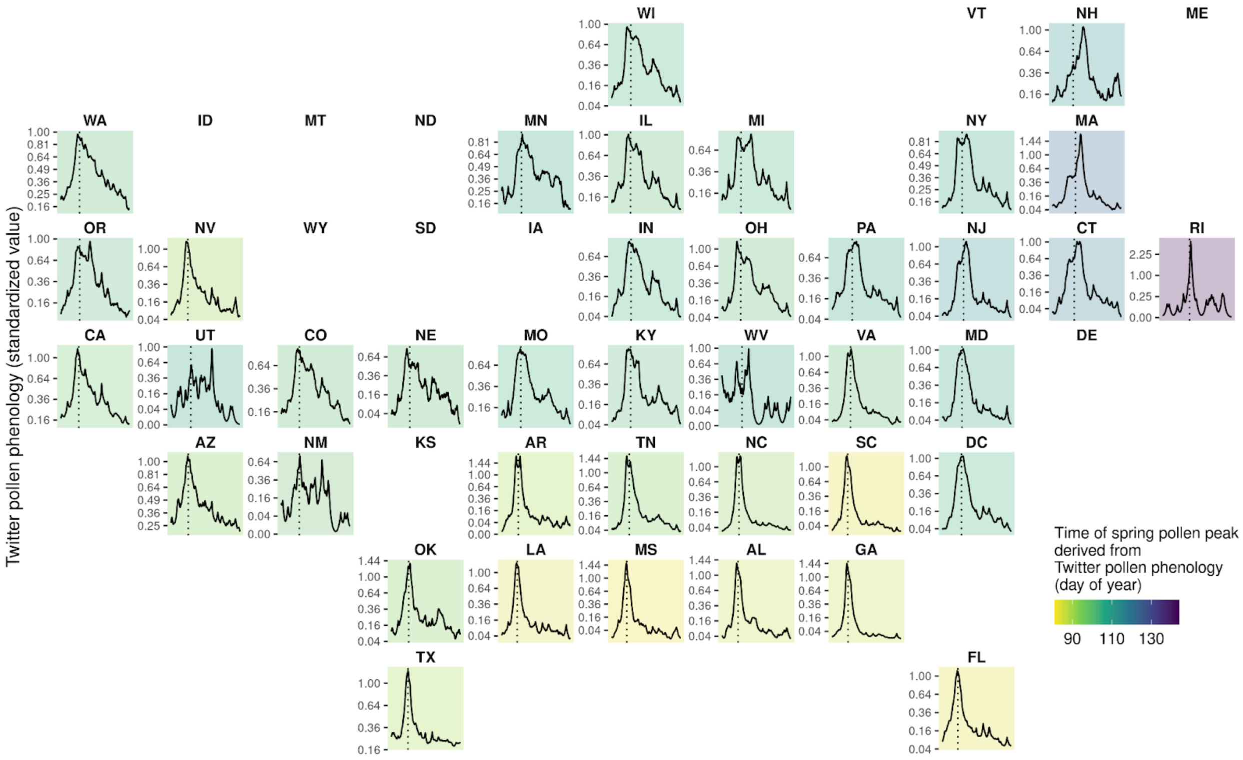


**Fig. S10.** Lines show the long-term average state-level Twitter pollen phenology across 39 states in the US, derived from the count of pollen-related tweets in each state. Dotted lines and colors of panels show the time of spring pollen peak derived from Twitter pollen phenology (day of year). A brighter color of the panel indicates an earlier spring pollen peak in the state.

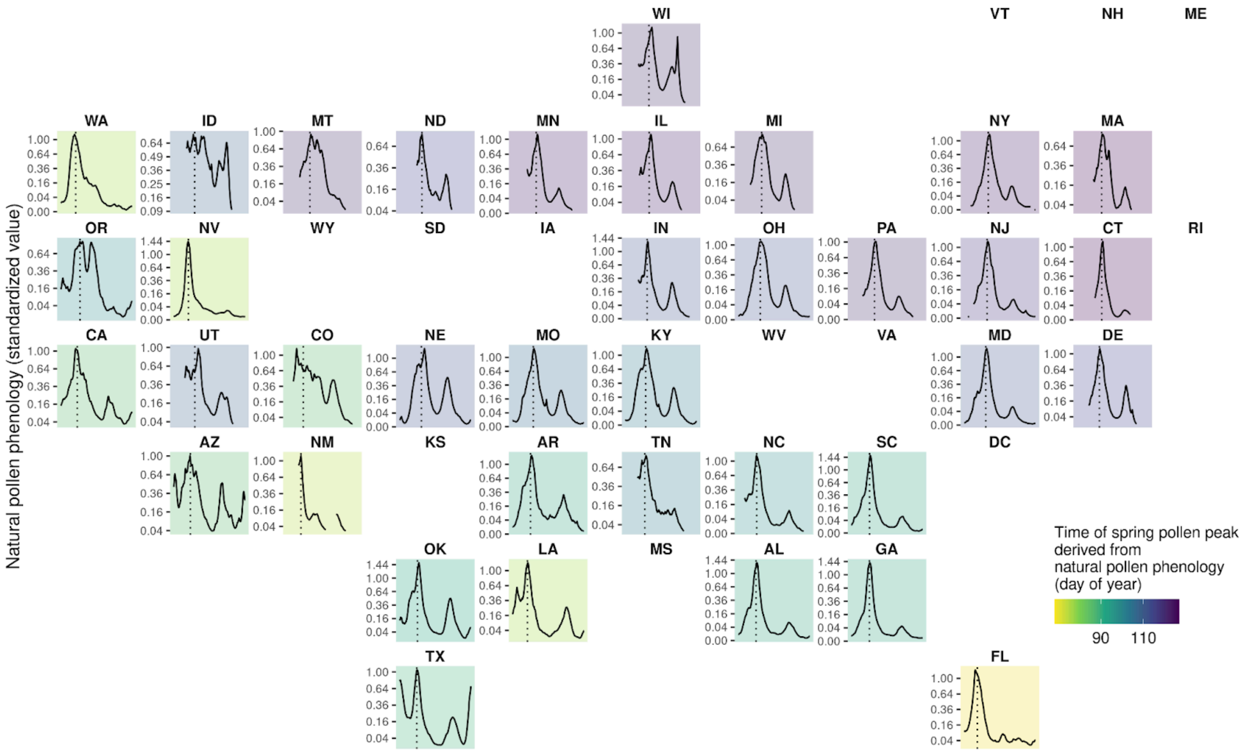


**Fig. S11.** Lines show the long-term average state-level natural pollen phenology across 37 states in the US, derived from the pollen concentration averaged from pollen counting stations in each state. Dotted lines and colors of panels show the time of spring pollen peak derived from natural pollen phenology (day of year). A brighter color of the panel indicates an earlier spring pollen peak in the state.

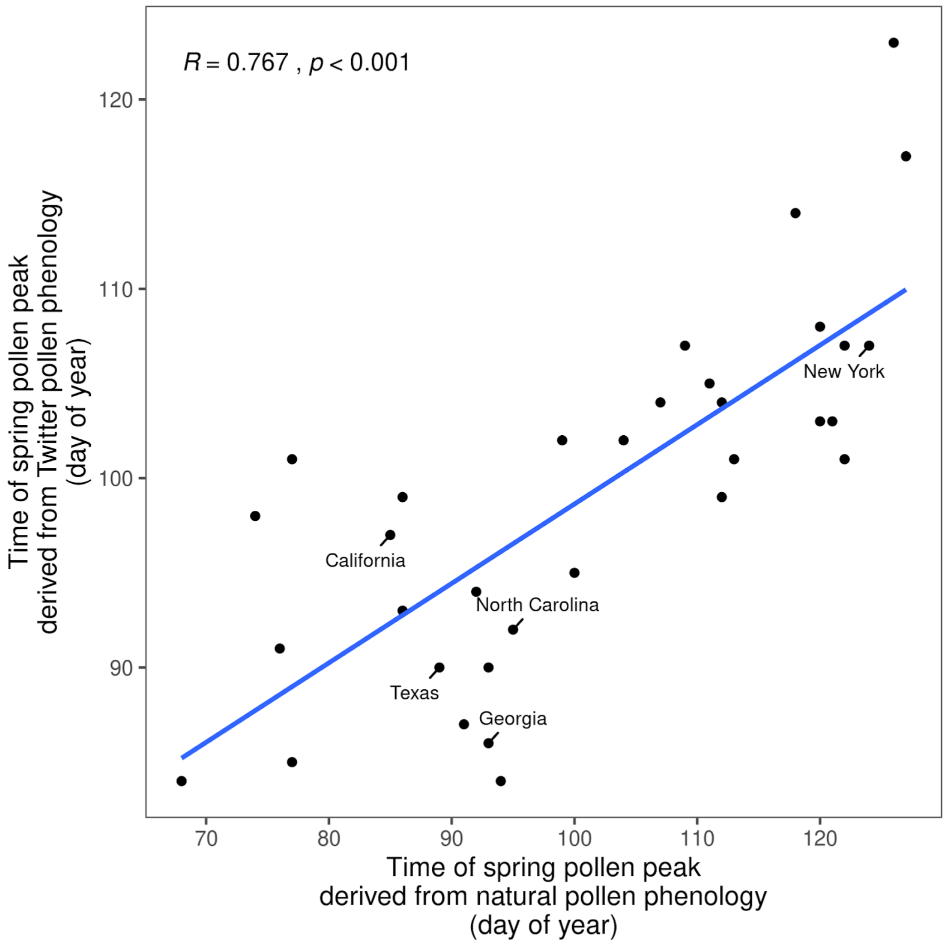


**Fig. S12.** Correlation between the time of spring pollen peak derived from Twitter pollen phenology and that derived from natural pollen phenology on the state level. Data points corresponding to selected states are labeled. Trend lines show the correlation across all states. Pearson correlation coefficient ($R$) and $p$-value ($t$-test) are shown.

### Supplemental Note

**Note S1.** Criteria for tweets in three nested groups

A) "pollen group" (e.g., "*this pollen is killing me*")

1) The tweet was about airborne pollen, i.e., pollen grains dispersed through the air. Discussions about pollen in fossils, pollen as human food, or pollen carried by pollinators, for example, did not qualify.

2) The tweet was time-sensitive. General statements about pollen allergies (e.g., "*I am allergic to dust and pollen*") and air filter commercials did not qualify.

B) "pollen-temperature group" (e.g., "*pollen level is high and warm weather will make it worse*")

1) The tweet satisfied all criteria in the "pollen group."

2) Temperature-related keywords were used in a context of temperature. Keywords when used in other contexts (e.g., "*my throat feels warm and itchy because of pollen allergy*", "*I am a hot mess in this pollen season*") did not qualify.

3) The tweet implied a short-term change in temperature (within a year or a few years). In particular, discussions on climate change did not qualify for this group.

4) The tweet implied correlation between temperature and pollen. Simply having both ideas in the same sentence without correlation (e.g., "*I can't go out because it is too hot and there is too much pollen*") did not qualify. Implications that warm temperatures and high pollen level usually co-occur (e.g., "*I wish I could have this warm weather all year minus the pollen*", "*spring brings warm weather but also pollen*") did qualify.

C) "pollen-climate group" (e.g., "*global warming could make your pollen allergies a lot worse*")

1) The tweet satisfied all criteria in the "pollen group."

2) The tweet implied a long-term change in temperature (on a decadal scale or longer). In particular, discussions on weather change did not qualify for this group.

3) The tweet implied causal impacts of climate change on pollen. Simply having both ideas in the same sentence without causation (i.e., "*there is global warming, wars, and a very high pollen count*"). We did not include tweets that discussed the effects of pollen on climate (i.e., "*find out about how pollen affects cloud formation and maybe climate*"), the effects of extreme heat reducing pollen activity (e.g., "*climate change = no pollen = good*"), or the understanding of pollen directly causing climate change (e.g., "*tree pollen causing climate change*").

4) The user did not disagree with the climate change impact on pollen phenology. We excluded tweets that explicitly expressed disagreement or skepticism (e.g, "*increase in pollen season caused by global warming???????*") or sarcasm (e.g., "*guess WaPo is going to recycle their old story about climate change making pollen season worse*").
